# Applying SCA for high-accuracy cortical auditory ERPs in children

**DOI:** 10.1101/2020.09.25.313809

**Authors:** S.E.P. Bruzzone, N. T. Haumann, M. Kliuchko, P. Vuust, E. Brattico

## Abstract

Overlapping neurophysiological signals are the main obstacle preventing from using cortical event-related potentials (ERPs) in clinical settings. Children ERPs are particularly affected by this problem, as their cerebral cortex is still maturing. To overcome this problem, we applied a new version of Spike-density Component Analysis (SCA), an analysis method recently introduced, to isolate with high accuracy the neural components of auditory ERP responses (AEPs) in 8-year-old children. Electroencephalography was used with 33 children to record AEPs to auditory stimuli varying in spectrotemporal features. Three different analysis approaches were adopted: the standard ERP analysis procedure, SCA with template-match (SCA-TM), and SCA with half-split average consistency (SCA-HSAC). SCA-HSAC most successfully allowed the extraction of AEPs for each child, revealing that the most consistent components were P1 and N2. An immature N1 component was also detected.

Superior accuracy in isolating neural components at the individual level even in children was demonstrated for SCA-HSAC over other SCA approaches. Reliable methods of extraction of neurophysiological signals at the individual level are crucial for the application of cortical AEPs for routine diagnostic exams in clinical settings both in children and adults.

**Highlights:**

- Spike-density component analysis (SCA) was validated on children ERPs
- SCA extracted overlapping neural components from auditory ERPs (AEPs)
- Child AEPs were modelled at the individual level

## Introduction

Auditory event-related potentials (AEPs) reflect changes in brain activity in response to auditory stimuli such as clicks, tones, or speech sounds. The earliest responses to auditory stimuli originate in the brainstem structures within the first 10ms from the stimulus onset and are referred to as brainstem auditory evoked responses (BAERs). In turn, longer latency components originate from cortical areas and are generally referred to as cortical auditory ERPs (CAEPs or AEPs) (Oken & Phillips, 2009). Over the past few decades, research has shown that CAEPs reflect cortical processes attributable to perceptual and cognitive functions, and changes in their amplitude and latency can reflect sensory deficits (Cone-wesson & Wunderlich, 2003; Eggermont & Ponton, 2002; Oken & Phillips, 2009) as well as cognitive impairments (Akshoomoff & Courchesne, 1994; Baldeweg et al., 2004; Čeponienė et al., 2009; Friedman, 2003; R Näätänen et al., 2012; Sunohara et al., 1999; Wiersema et al., 2005). Furthermore, several studies demonstrated CAEP as putative indices of experience-dependent plasticity in learning processes (Lappe et al., 2008; Näätänen, 2009; Shahin et al., 2003, 2004; Trainor et al., 2011) and auditory recovery following cochlear implantation (Mehta et al., 2019; Petersen et al., 2020; Sharma et al., 2002, 2015). However, there are certain limitations attributed to the nature of CAEPs that currently constrain their usage in clinical settings, enabling only BAERs to be adopted in clinical practice.

The main limitation of CAEPs is represented by the difficulty in achieving reliable single neural responses at the individual level. Such high inter-subject variability is mostly imputable to individual differences in brain structures and to each CAEP being the summation of a series of components originating by different cortical sources and overlapping over time (Čeponienė et al., 2005; Picton et al., 1974; Ruhnau et al., 2011; Sussman et al., 2008). Consequently, cortical evoked responses are easily masked by other interfering signals, such as earlier or subsequent components and strong endogenous oscillations such as alpha waves. These phenomena challenge the isolation of single CAEP components, causing inaccurate approximations and unreliable estimations (Oken & Phillips, 2009; Scharf & Nestler, 2018) that lead to low CAEPs replication rates at the single-subject level (Luck & Gaspelin, 2017).

To tackle this problem, the recent study by Haumann and colleagues (Haumann et al., 2020) utilized spike-density component analysis (SCA), a novel method that allows us to isolate overlapping neural components from magneto- and electroencephalography (MEG and EEG) signals. SCA is based on the principle that the same stochastic pattern observed from intracranial recording in single neurons and larger neuronal assemblies (Maimon & Assad, 2009; Stein et al., 2005) is reflected in the larger-scale cortical activity measured with M/EEG (Shin, 2002). Under this assumption, it models the spatial topography, the polarity, and the temporal shape of single neural components by means of Gaussian probability density functions (Beauducel, 2018). The analysis procedure involves decomposing the individual average waveforms into their constituent spatiotemporal components. SCA decomposition provides a more accurate representation of AEPs in adults than decompositions performed with independent component analysis (ICA) and principal component analysis (PCA) or extractions done with conventional averaging methods (Haumann et al. 2020).

The first results obtained with SCA have demonstrated its high accuracy in extracting neural components of interest from both MEG and EEG signals recorded with adults. The component of interest for every adult subject was automatically selected from the SCA decomposition using a template matching procedure in which each individual ERP waveform was compared to the group-level average ERP and, if matching, the component was extracted (Haumann et al. 2020).

However, an additional challenge arises when studying a developing brain in a children population that is not characterized with the same structural and functional organization as an already matured brain of an adult (Bishop et al., 2007; Čeponiene et al., 2002; Lippé et al., 2009; Picton et al., 1974; Ruhnau et al., 2011; Tonnquist-Uhlén et al., 1995).

Ongoing maturational processes occurring across development affect the neural responses recorded at the scalp (Wunderlich & Cone-Wesson, 2006). While the obligatory cortical responses in adults are constituted by the P1-N1-P2 complex (occurring in within 50ms-300ms), in infants and children they are represented by P1 and N2 (Albrecht et al., 2000; Caviness et al., 1996; Snook et al., 2005; Tonnquist-Uhlén et al., 1995). Before 10 years of age, N1/P2 emerge only in response to stimuli presented with interstimulus intervals (ISIs) longer than 1s. N1 amplitude progressively increases throughout development and the ISI length required for its appearance gradually decreases with age, whereas P1 and N2 amplitudes decrease (Čeponiene et al., 2002; Kushnerenko et al., 2002; Sussman et al., 2008). Moreover, changes in amplitude are accompanied by a general shortening of the CAEPs latency (Bishop et al., 2007; Čeponiene et al., 2002; Habibi et al., 2016; Ruhnau et al., 2011; Wunderlich & Cone-Wesson, 2006). Therefore, it has been suggested that N1 components, although already present in early stages of life, might be masked by the most prominent P1 and N2 until adolescence (Čeponiene et al., 2002, 2005; Paetau et al., 1995; Sharma et al., 1997) and only stabilize in adulthood (Ponton et al., 2000). In addition, children’s neural responses are less stable and less homogeneous among individuals of the same age, due to different developmental rates (McIntosh et al., 2014; Mueller et al., 2008; Snyder et al., 2002).

Therefore, in the case of children, constraining the analyses by comparing individual responses to the average group signal does not represent the optimal strategy in this case, as it assumes that the latency values and scalp topographies are consistent across all participants. Hence, in this study, we complement the standard SCA procedure (SCA template matching or SCA-TM), with an additional approach, namely the half-split average consistency (Carter et al., 2010) applied to SCA (SCA-HSAC). SCA-HSAC extracts neural components from the individual waveforms by searching for the most consistent neural components in the SCA decomposition of each subject’s averaged data, thus enabling components extractions at the single-subject level. This allows us to model neural responses with higher inter- and intra-subject reliability

In this study, we tested this approach on the data from 8-year old children, whose obligatory CAEPs were recorded with EEG. The CAEPs were elicited by auditory stimuli differing by their spectrotemporal properties, consisting of sounds played on three different instruments: piano, flute, and violin. Differences in acoustic features such as spectral properties are known to be reflected in changes of adult AEPs, both in normal and impaired hearing adults (Jones et al., 1998; Seol et al., 2011; Shahin et al., 2005). We, therefore, hypothesized that SCA would also highlight the differences in response to stimuli characterized by distinct spectrotemporal features.

In summary, the present study aimed to: 1) validate that child EEG signals (affected by the maturational changes occurring in early life) can be decomposed into SCA components and that child’s AEPs can be identified from the SCA decompositions, 2) model neurophysiological differences in children responses to the spectral properties of sound with SCA. The main hypotheses in this regard were: a) SCA-HSAC would provide similar or more accurate results to SCA with template matching and conventional children CAEP waveforms (containing component mixtures), b) SCA would retain or increase effect sizes for effects of different instrument sounds on ERP amplitudes, when compared to the original ERP amplitudes.

## Material and methods

### Participants

We analysed data recorded from 33 eight-year-old (99.8±4.5 months) children, 18 females and 15 males from a pool of 40 children, recruited from the second-grade pupils of a public school in Silkeborg Kommune, Denmark. All participants were in normal health condition and had normal hearing according to the information provided by their parents who filled the children and family background questionnaire (Kliuchko, 2017; Müllensiefen et al., 2014; Ukkola-Vuoti et al., 2013) Originally, informed written consent for the study was received from parents of 40 children. The oral information about the study participation was also explained to each child prior to a measurement. One child did not give assent to participate in the study, two children withdrew during the preparation or recording, and one child was not in the school on the measurement day. Data from three of the subjects were excluded from the final dataset due to bad quality. The research protocol was approved by the Institutional Review Board (case number DNC-IRB-2019-004) and was conducted in accordance with the principles of the Declaration of Helsinki. Information about the study and participation invitation were distributed to all parents via the school’s intranet.

### Stimuli and Procedure

The stimuli in this study consisted of tones with three different timbres (piano, flute and violin) and two frequencies, corresponding to musical pitch F and C of the 4^th^ and 5^th^ octave respectively. Piano tones were generated using the sample sounds of Wizoo acoustic piano from the software ‘Alicia’s Keys’ in Cubase (Steinberg Media Technologies GmBH). Flute and violin tones were created by transforming the timbre of the piano sounds on Adobe Audition (Adobe Systems Incorporated). All sounds were normalized. Each tone had a duration of 300ms (5ms rise and fall) and the presentation order of the three stimuli types were randomized. The tones were separated by an interstimulus interval (ISI) of variable duration 2s (± 0.5s). A total of 144 tones (trials) were presented. The three conditions were evenly distributed across the stimulation. Before the above-mentioned stimulation, participants were presented with a no-standard musical multifeature MMN paradigm (Kliuchko et al., 2016) in two blocks with the total duration of approximately 14 minutes. These data will be reported in a separate paper.

The sound stimuli were presented with the presentation software (Neurobehavioral Systems, Albany, USA) through headphones. The loudness of the stimuli was set constant for all subjects. Prior to the measurement, a soundcheck was done to assure that the sound level was comfortable for each participant. All measurements were carried out in the premises of the school on the same day. Participants sat in a chair in the middle of the room, in front of a table with a laptop that played cartoons or kids shows of the child’s personal choice. During the preparation, the show was played with sound, which was then switched off during the measurement. The participants were instructed to watch the cartoons and ignore the sounds in their headphones. They were also asked to sit as still as possible during the recording. A neck pillow and a small stool to place feet on were used to further reduce potential movements. Researchers were out of sight to a participant but present in the room during the recording.

### EEG data acquisition and preprocessing

Brain activity was recorded using a mobile EEG setup and a 32 channels cap (EasyCap, actiCap) with Ag-AgCl electrodes. Eye movements and blinks were tracked by placing electrooculography (EOG) electrodes on the external eye corners, above the left eyebrow, and on the cheek below the right eye. An additional electrode was placed on the nose and used as an offline reference. The channel used as an online reference was FCz. EEG signals were taken with a 1000Hz sampling rate.

EEG data were first analyzed with Matlab-based opensource toolbox EEGLAB (Delorme & Makeig, 2004) and with the ERPlab plugin (Lopez-Calderon & Luck, 2014). Raw data were re-referenced offline to the average of the left and right mastoids and downsampled to 500Hz. After filtering with a 1Hz high-pass filter and a 30Hz low-pass filter, the EEG signals were notch-filtered at 45-55Hz with the CleanLine plugin (Mullen, 2012), to filter out line frequency noise. The data were inspected by eye and a maximum of three noisy channels were removed. Independent component analysis (ICA) was then applied to remove eye artefacts and a maximum of four artefactual components was rejected. After ICA, the removed channels were interpolated. 500ms epochs (100ms pre-stimulus and 400ms post-stimulus), time-locked to the presentation of each tone, were created. Epochs were removed if they involved an amplitude change exceeding a threshold of ±100μV.

Three different methods were used to obtain the final ERP signal, giving rise to three different average waveforms. An “original waveform” was obtained by averaging the epoched files. The other two waveforms were obtained by applying SCA decomposition on the averaged epoched data: the SCA components of interest were isolated individually. In particular, SCA components were extracted with two different approaches: one approach involved matching the individually extracted components to the grand average waveform, obtained by the average across all subjects (SCA with template match) (Haumann et al., 2020); the other approach instead, involved extracting the components that were most consistently present across half of the trials from the individual waveforms (SCA with half-split average consistency). The two SCA-based methods gave rise to the “SCA-TM” and “SCA-HSAC” waveforms respectively.

### Spike density component analysis (SCA)

The spike density component analysis (SCA) method (Haumann et al., 2020) is an open-source, Field-Trip-compatible (version r9093, Donders Institute for Brain, Cognition and Behaviour/Max Planck Institute, Nijmegen, the Netherlands) Matlab (MathWorks, Natick, Massachusetts) function. SCA allows to isolate neural sources with high temporal and spatial resolution, by modelling their spatial topography, polarity and temporal shape with temporal Gaussian functions. The SCA function is applied to the average individual waveforms (the epoched files) with the following assumptions: a) EEG waveforms in the time domain can be modelled with a Gaussian function; b) components have a signal-to-noise and interference ratios SNIR>1; c) components differ in time, width across time or topography.

The analysis proceeds as follows. First, the SCA function finds the maximum amplitude across channels and time. The component waveform is modelled by estimating the Gaussian function parameters and fitting it to the signal. Then, the component weighting matrix is estimated by means of linear regression and multiplied by the channel weight vector. Finally, the component waveform is subtracted from the multichannel waveforms, and residual waveforms are obtained. This operation is repeated iteratively, based on minimizing the sum of the residual waveforms across channels and time. The resulting file contains all the overlapping components that have been modelled individually, each one with its spatial topography and temporal morphology.

### Template matching (TM)

Once that the EEG waveforms have been decomposed with SCA at the individual level, the SCA components of interest (reflecting P1 and N2 in this case) are isolated from the rest of the EEG signal. The previously validated SCA pipeline (referred to as SCA-TM) implied the extraction of the component of interest of each subject by matching individual components with the grand average waveform across all subjects. This is done by means of an automatized method that involves comparing the component with the average waveform and extracting it when the two signals match (Haumann et al., 2020). A weakness of template matching is that it requires the evoked responses to be morphologically homogenous within groups, i.e., similar in latency and width within a group. This is expected to be problematic in relation to children that undergo relatively large changes in ERP morphology related to brain maturation processes.

### Half-split average consistency (HSAC)

Given the high latency differences across child subjects in this study, we also adopted a half-split average consistency procedure, an improvement to the former approach that allows to extract the most consistent components from the individual average waveforms. Carter et al. (2010) suggested a visual inspection of signal reliability in clinical procedures. This involved dividing half-split averages into odd and even numbered trials and detecting whether the AEP of interest is visible or not. Adding more half-split averages allows more reliable statistical inference of the signal consistency. However, a large total number of trial combinations is possible: e.g., with just a small sample of only 10 trials there is already 10!/(10/2)!=30240 total possible combinations of half-split averages. The half-split average consistency approach (referred to as SCA-HSAC) solves this problem by repeatedly taking half of the total number of trials in a randomized manner with an equal chance of drawing each trial from a uniform distribution (by means of a Monte Carlo simulation) and finding the most consistent components across the half-split averages (HSAs).

Components were considered as consistent when they were found in 70% of the half-split averages. The half-split average consistency was tested for each component. The testing procedure involved subtracting all the components other than the tested one from the HSAs. Next, the HSAs were transformed into component space using the sum of the HSAs across EEG channels weighted by the channel weights for the tested SCA component:

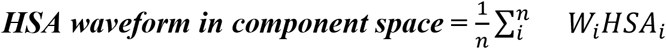

(where *i* is the channel number, *W* is the channel weight for the SCA component, HSA is the half-split average waveform).

Further, the computational processing speed and accuracy of the component identification was increased by initially constraining the analysis to specific channels, latencies and polarities prior to the HSAC testing. In this case, a region of interest (ROI) constraint was applied for P1 extraction by searching for positive peaks in one of 12 frontal channels (*F3, Fz, F4, FC1, FC2, FC5, FC6, F7, F8, C3, Cz, C4*), whereas N2 extraction included a ROI with negative peaks over 20 channels (*F3,Fz,F4,FC1, Fp1, Fp2, FC5, FC6, F7, F8, T7, T8, CP1, CP2, CP5, CP6, FC2,C3,Cz,C4*). Latency ranges for component identification were 0-130ms for P1 and 200-350ms for N2. The choice of the channels and latencies was made on the basis of the channels showing the strongest amplitudes across participants for the component of interest while allowing variability across the individual children.

The pipeline of the SCA decomposition is illustrated in Figure 1. We compared the efficiency of the conventional method for ERP extraction with the SCA template match (SCA-TM) and half-split average consistency approaches (SCA-HSAC).

**Figure 1.**
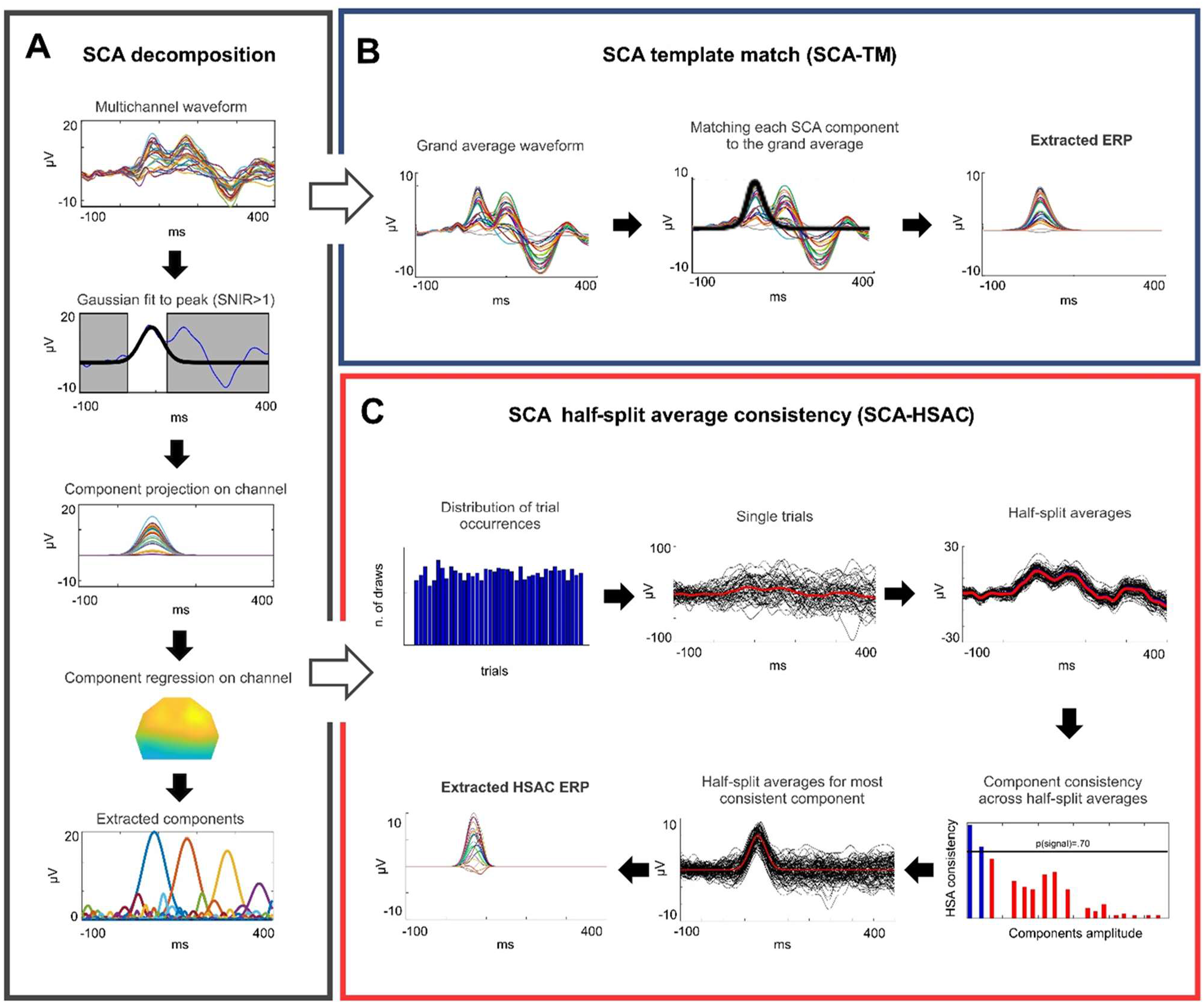
Overview of the spike density component analysis (SCA) pipeline. The original waveform is decomposed into its components by fitting Gaussian temporal functions and projecting the component signal to its topography (A). For every subject, the components of interest are extracted if they match the grand average waveform, with the template matching approach (B), or by searching for the most consistent components across half of the trials with the half-split average consistency approach (C).

### ERP Analyses

Amplitude values were identified automatically within a time window of 50ms (for P1) and 30ms (for N2) around the maximum peak of the grand average waveform for each condition. Mean amplitude values were calculated as the average value across the same channels (12 frontal channels for P1 and 20 frontal and frontoposterior channels for N2) that were used in the SCA-HSAC extraction. Latency values were identified automatically by searching the maximum peak within a 0-130ms time window for P1, 130-200 for P2 and 200-350ms for N2. Time constraints were chosen by visual inspection of the individual waveforms across all conditions.

### Statistical analyses

Statistical analyses were performed on MATLAB. The normality of the data distribution was assessed visually (*hist* function) and with the Kolmogorov-Smirnov test for normal distribution (*kstest* function). As the data were not normally distributed, non-parametric tests were performed. Friedman’s ANOVA (*friedman* function) was used to test differences among repeated measures across the three conditions (Table 2). The Wilcoxon signed-rank test (*signrank* function) was used as a paired-test and the relative *r* value as a measure of an effect size of spectral sound differences on P1 and N2 (Table 3). Analyses were conducted on the three average waveforms: the original waveform, the SCA-HSAC waveform and the SCA-TM.

## Results

Median and interquartile range values for P1 and N2 amplitude and latency are reported in Table 1; results from Friedman’s ANOVA in Table 2 and results from the Wilcoxon-signed rank test for the amplitudes and latencies in Table 3.

**Table 1.** Median and respective interquartile range (IQR) of amplitude and latency values for each condition.

| ER<br>P | Condition | Amplitude (μV) | | | | | | Latency (ms) | | | | | |
| --- | --- | --- | --- | --- | --- | --- | --- | --- | --- | --- | --- | --- | --- |
|  |  | Original | IQ<br>R | SCA-HSAC | IQR | SCA-TM | IQR | Original | IQ<br>R | SCA-HSAC | IQ<br>R | SCA-TM | IQR |
| P1 | Piano | 6.84 | 3.88 | 6.99 | 4.04 | 6.85 | 4.65 | 83 | 6 | 86 | 12 | 82 | 15 |
|  | Flute | 8.59 | 2.8 | 9.17 | 3.39 | 8.35 | 5.38 | 94 | 14 | 96 | 24 | 92 | 18 |
|  | Violin | 7.82 | 3.6 | 7.64 | 4.37 | 7.53 | 5.58 | 92 | 16 | 94 | 30 | 88 | 26 |
| N2 | Piano | -6.64 | 3.35 | -7.11 | 3.75 | -6.98 | 3.88 | 259 | 21 | 253 | 21 | 254 | 18 |
|  | Flute | -4.08 | 2.77 | -3.79 | 3.19 | -3.94 | 3.38 | 262 | 32 | 262 | 22 | 268 | 22 |
|  | Violin | -3.34 | 2.83 | -3.27 | 3.46 | -2.16 | 3.71 | 270 | 39 | 260 | 39 | 252 | 21 |

**Table 2.** Results from Friedman’s ANOVA: degrees of freedom (dF), p-value (P) and chi-squared (χ^2^).

|  |  | Amplitude |  |  | Latency |  |  |
| --- | --- | --- | --- | --- | --- | --- | --- |
| ERP | waveform | dF | P | $\chi^2$ | dF | P | $\chi^2$ |
| P1 | Original | 98 | <.001 | 20.06 | 98 | <.001 | 19.44 |
|  | SCA-HSAC | 98 | <.001 | 14.97 | 98 | <.01 | 11.10 |
|  | SCA-TM | 98 | <.01 | 9.66 | 98 | <.001 | 18.06 |
| N2 | Original | 98 | <.001 | 22.06 | 98 | <.05 | 6.73 |
|  | SCA-HSAC | 98 | <.001 | 25.88 | 98 | .144 | 3.88 |
|  | SCA-TM | 98 | <.001 | 35.09 | 98 | <.001 | 13.15 |

**Table 3.**
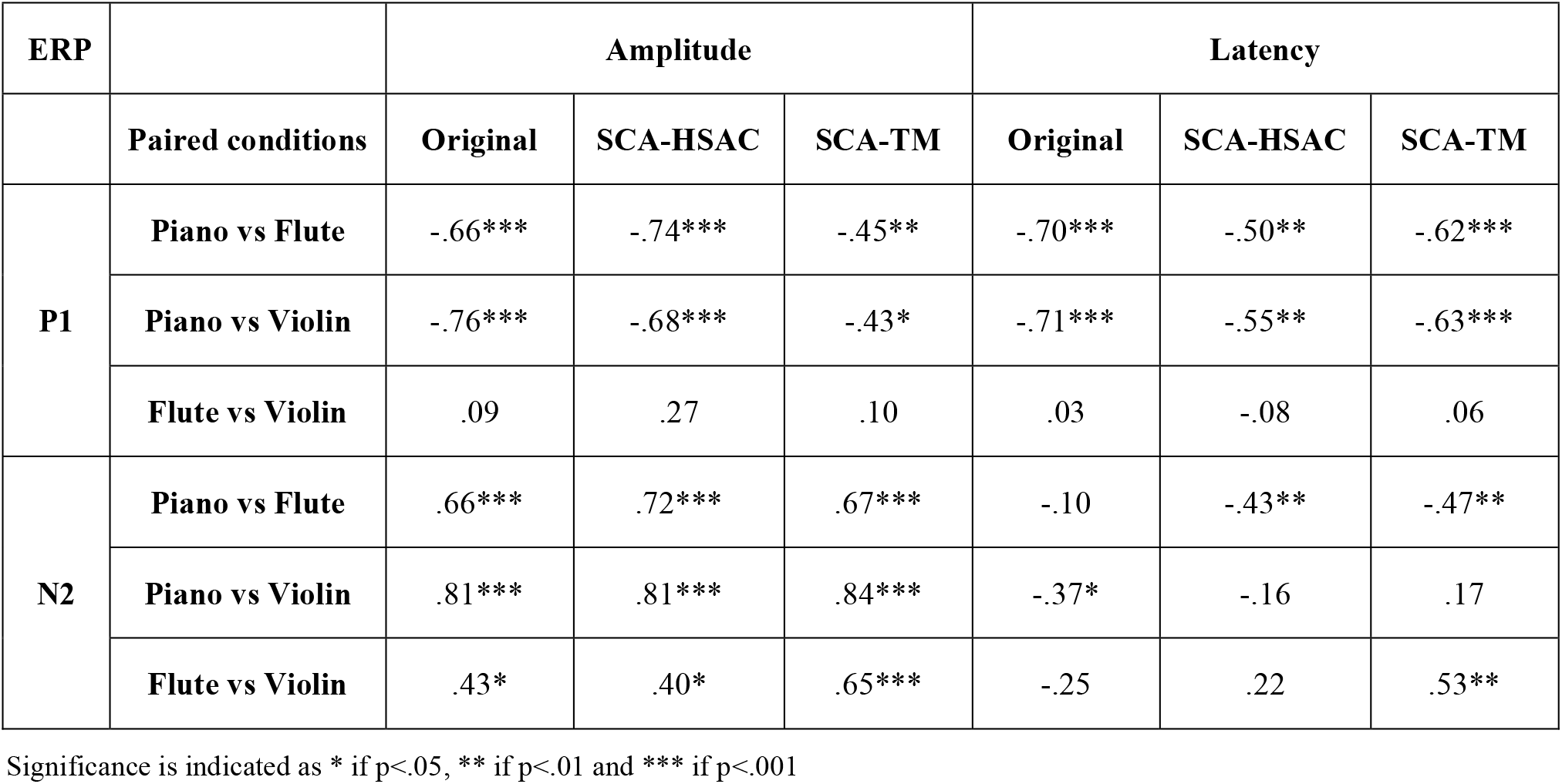
r values for effect size from the Wilcoxon-signed rank test. The difference between the conditions is moderate when 0.3<r<.5, large when r>0.5

### Explained variance with SCA

SCA decomposition successfully modelled an average of 178 SCA components per child for the piano condition, 184 for the flute, and 189 components for the violin condition. The explained variance of the total signal by the SCA components was equal to 99.996% for each condition.

### SCA with half-split average consistency

As expected, the half-split average procedure successfully extracted P1 and N2 for all children, confirming that the most consistent components at this developmental stage are P1 and N2 (Figure 2). Furthermore, SCA-HSAC allowed extracting P2 components, although not clearly identifiable from the grand average waveforms. However, P2 was not as consistent as P1/N2 across the subjects: P2 was extracted for 29/33 subjects in the piano conditions, 30/33 in the flute condition, and 28/33 in the violin condition. Therefore, only P1 and N2 will be discussed more in detail.

**Figure 2.**
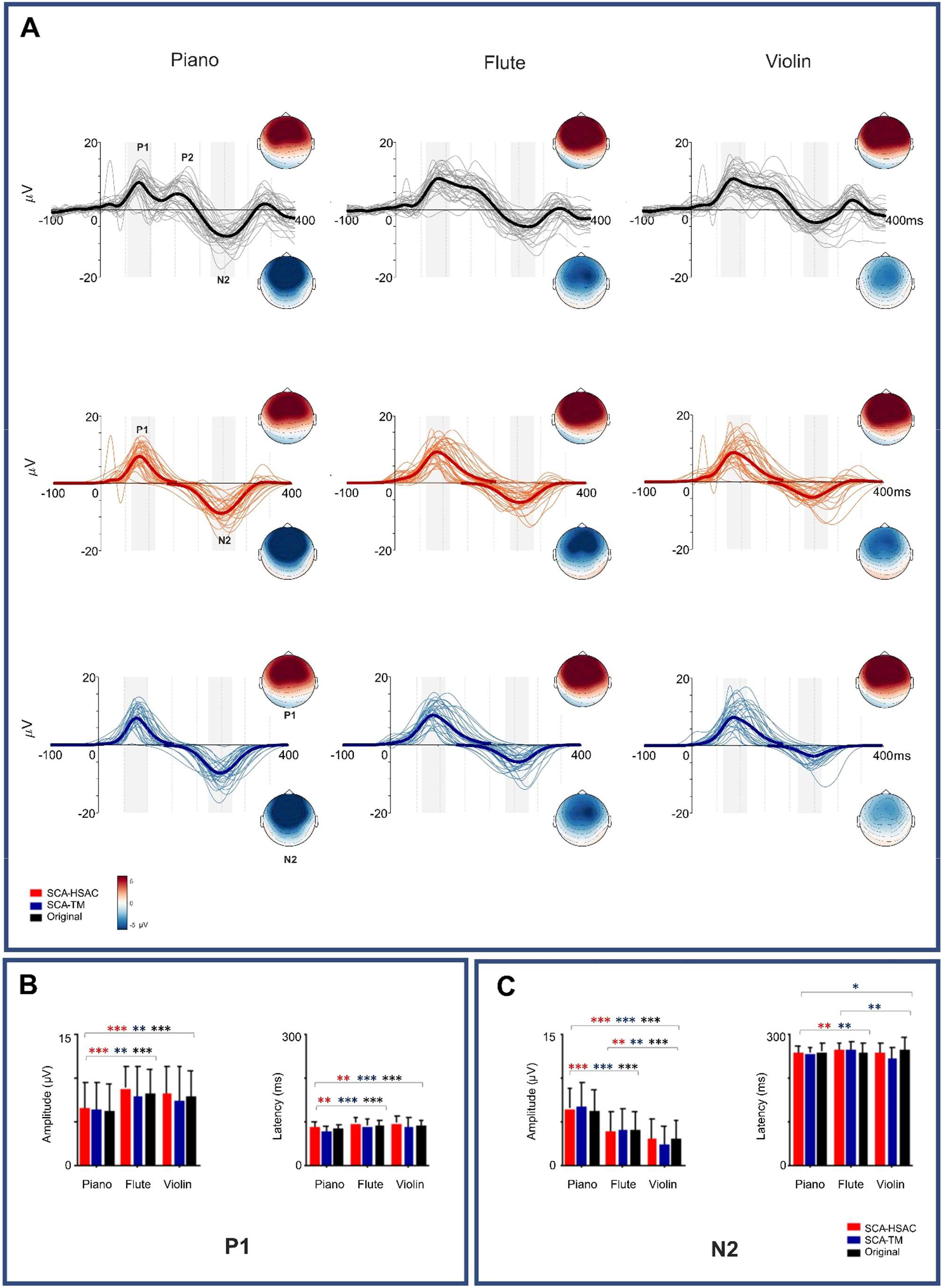
Original (black), SCA-HSAC (red) and SCA-TM (blue) waveforms of P1 and N2 and relative topographies for piano, flute and violin conditions (A). The thicker lines represent the grand average across the channels considered for the statistical analyses, whereas the thinner lines represent the individual waveforms of single subjects. Mean amplitude and latency values of each condition in the three methods for P1 (B) and N2 (C). Significance levels are indicated by * p<.05, ** p<.01 and *** p<.001.

P1 and N2 topography maps showed their characteristic peak at frontocentral electrodes. Waveforms and topoplots for P1 and N2 in the three conditions are illustrated in Figure 2. Moreover, visual inspection of the components extracted with SCA also revealed an N1-like response (Figure 3), peaking around 90ms-150ms. However, neither SCA-HSAC nor SCA-TM allowed to extract it from each subject.

**Figure 3.**
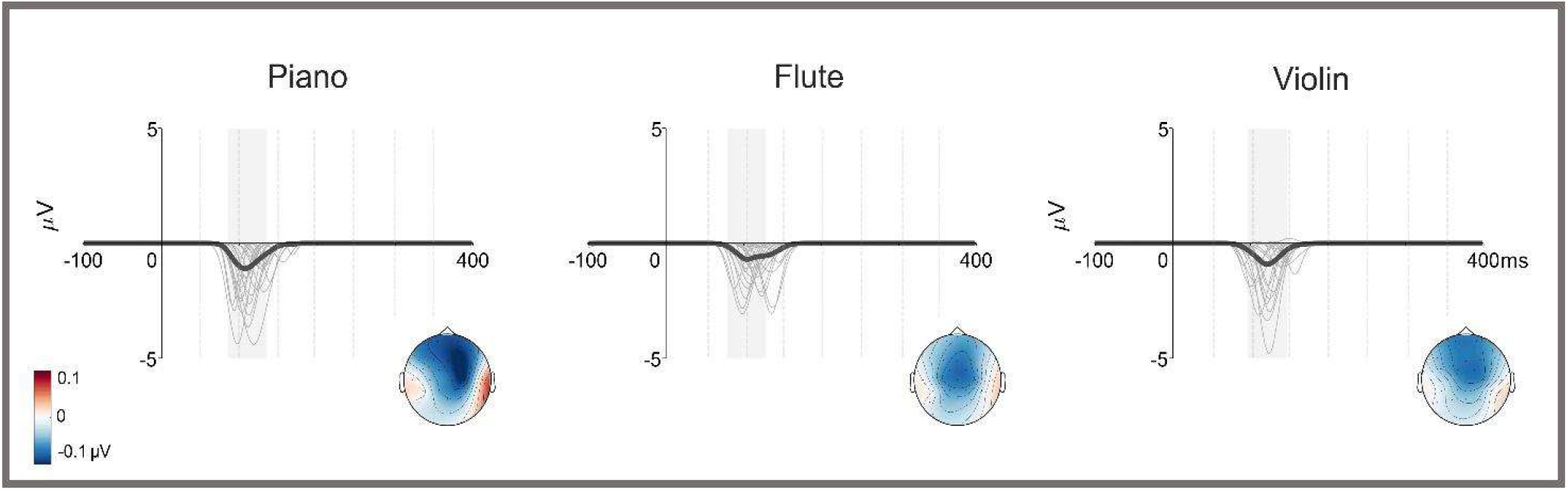
Manual inspection of the components extracted by SCA revealed an N1-like negativity, although not clearly detectable from the grand average waveforms. Here the grand average and individual waveforms with relative topography for the N1-like component in piano, flute and violin conditions.

### SCA with template matching

The template match procedure only successfully matched the components to the grand average waveform for both P1 and N2 and across all conditions for 17/33 participants. The individual P1 components were not matched for 2/33 subjects in every condition (with the subjects without a match being different in every condition). The individual N2 responses instead, were not matched for 2/33 subjects in the piano condition, for 5/33 in the flute condition and 12/33 in the violin condition. Thus, as expected, the TM method failed to identify more of the child ERPs, although, they were visible in the original ERP waveforms.

### Effects of instrument sound on ERPs

#### P1 amplitude

Friedman’s ANOVA on the original waveforms revealed significant differences across piano, flute, and violin conditions (χ^2^_F_(2)=20.06, P<.001). Paired-test analyses with Wilcoxon signed-rank test indicated a significant difference between piano and flute and between piano and violin, with the piano amplitude being considerably smaller than the flute and violin amplitude (Table 3). No significant differences were found between flute and violin conditions.

ANOVA on waveforms analysed with SCA-HSAC and SCA-TM confirmed the significant differences across conditions found in the original waveforms (SCA-HSAC: χ^2^_F_(2)=14.97, P<.001; SCA-TM: χ^2^_F_(2)= 9.66, P<.001). Piano amplitude was significantly smaller than the flute condition both in SCA-HSAC and in SCA-TM, although SCA-HSAC revealed greater differences between the two conditions, closer than SCA-TM to the original value (Table 3). Analogously, a significant difference was found between piano and violin conditions, with a greater effect for the SCA-HSAC waveform compared to the SCA-TM one (Table 3). No significant differences were found between flute and violin conditions.

Effect sizes on the SCA-TM waveform were lower than both those of the SCA-HSAC and those of the original waveform (Table 3), probably due to the lower number of subjects for whom the template match was successful.

### P1 latency

Significant differences across the three conditions were also found in P1 latency (χ^2^_F_(2)=19.44, P<.001). Similar to P1 amplitude, piano latency was significantly shorter than those of both flute and violin (Table 3). The contrast between flute and violin instead, did not indicate significant differences.

As visible in Table 3, similar findings from the Friedman’s test were found in SCA-HSAC (χ^2^_F_(2)=11.10, P<.005) and SCA-TM (χ^2^_F_(2)=18.06, P<.001) waveforms. Wilcoxon-signed rank showed a significantly shorter latency for piano in contrast to flute conditions. Similarly, the piano vs violin contrast revealed a significantly shorter latency for piano. In both contrasts, a greater difference between the two conditions was found in the original waveform and SCA-TM compared to the SCA-HSAC. No significant differences were found between flute and violin latencies.

### N2 amplitude

Significant differences across conditions were also found for N2 amplitude (χ^2^_F_(2)=20.06, P<0.001), whereas paired-test analyses on the piano vs flute and piano vs violin conditions showed a significantly larger amplitude for piano compared both to flute and violin (Table 3). Furthermore, the flute vs violin contrast revealed that flute amplitude was significantly bigger than that of violin, although the difference was smaller than in the other contrasts (Table 3).

Differences across conditions were significant also in SCA-HSAC (χ^2^_F_(2)=25.88, P<.001) and SCA-TM (χ^2^_F_(2)=35.09, P<.001) waveforms. Paired-test analyses confirmed the results of the original waveform: piano amplitude was significantly larger than flute and violin in both SCA-HSAC and SCA-TM waveforms as well as flute had a larger amplitude compared to violin, with a greater effect size for the SCA-TM waveform compared to the SCA-HSAC and the original waveforms (Table 3).

### N2 latency

Original N2 latency values were significantly different across conditions (χ^2^_F_(2)=6.73, P<.05). Piano vs violin contrast revealed a significantly shorter latency for the piano compared to the violin condition (Table 3). The piano vs flute and flute vs piano contrast instead, did not reveal significant differences.

Friedman’s test revealed significant differences across conditions for SCA-TM (χ^2^_F_(2)=13.15, P<.001) but not for SCA-HSAC waveforms (χ^2^_F_(2)=3.88, P=0.14). The piano vs flute contrast showed similar significantly shorter piano latencies compared to flute latencies in both SCA-HSAC and SCA-TM waveforms (Table 3). The piano vs violin contrast did not render significant differences in either waveform. Interestingly, a significant difference in SCA-TM but not in SCA-HSAC nor in the original waveform was found between flute and violin (Table 3). However, as the SCA-TM did not succeed in extracting the components for 12/33 subjects in the violin condition, such significance might be less reliable due to the smaller sample considered.

### Application of SCA to isolate P1 from interfering N1 component

Further visual inspection on the SCA components revealed a weak frontocentral negativity (maximum peaks at channels Fp1, Fp2, Fz, F3, F4, FC1, FC2, Cz, C4, CP2) with reversed polarity near the mastoids (Figure 3), as typical of components originating in the auditory cortex. The negative AEP occurred between the two main positive peaks (P1 and P2) or, if P2 was not present or not detectable, right after the first main positive peak, within the time window 90-160ms (piano: Mdn=114, IQR=24; flute: Mdn=114, IQR=30; violin: M=124, IQR=24). The peaking latency of such negative component often partially overlapped the earlier P1 in children, which might have caused confounding effects between P1 and N1 components. This might explain the longer latencies and increased between-subject variance in P1 latency following SCA-HSAC extraction compared to the original waveform, as well as the greater effect size for the original waveforms of the piano vs flute (r=-.70) and the piano vs violin (r=-.71) contrasts, compared to those obtained with the SCA methods (SCA-HSAC: r=-.50, r=-.55; SCA-TM: r=-.62, r=-.63). The median amplitudes were -3.10μV (IQR=2.80), -2.90μV (IQR=1.60), and -2.50μV (IQR=2.10) for the piano, flute, and violin respectively.

## Discussion

In this study we applied SCA for obtaining high-accuracy children’s AEP signals to sounds with different spectrotemporal characteristics.

Waveforms analyzed with SCA outperformed over those analyzed with conventional ERP analysis methods. Two different approaches were adopted for the SCA analysis: SCA-TM and SCA-HSAC. SCA-HSAC revealed to be the most accurate approach for the extraction of child AEPs, providing improved estimations of the brain signals at the individual level. The higher accuracy is reflected by the complete number of children for which AEPs were identified, the resemblance of the HSAC-SCA waveform features to those of the original waveforms (as indicated by the similar amplitude and the effect size values) and the ability of SCA-HSAC to isolate P1 from an N1-like overlapping component. Additionally, HSAC-SCA indicated P1 and N2 as the most consistent components across all subjects, in line with previous findings showing the predominance of P1 and N2 responses in early stages of brain development (Čeponienė et al., 2005; Paetau et al., 1995; Picton et al., 1974; Ruhnau et al., 2011; Sharma et al., 1997; Sussman et al., 2008). Conversely, SCA-TM revealed a limitation in extracting components from a sample with non-homogeneous latencies.

The main differences across methods were found in the latency values. Specifically, P1 latency with SCA-HSAC revealed smaller effect sizes for piano vs flute and piano vs violin contrasts, compared to the original and the SCA-TM waveform. Manual inspection of the components extracted with SCA revealed a negative frontocentral component between P1 and P2 with the typical latency (90-160ms) and frontocentral topography of N1 (Čeponiene et al., 2002; Näätänen & Picton, 1987; Sharma et al., 2015; Tonnquist-Uhlén et al., 1995; Wunderlich & Cone-Wesson, 2006). It has previously been suggested that shorter latency of P1 in original EEG waveforms might be due to the fusion of the N1 component with P1, which can lead to overestimating the effects attributed to P1 amplitude and latency (Čeponiene et al. 2002). Following SCA decomposition, P1 amplitude and latency increased, most likely due to the subtraction of the overlapping N1. Likewise, the greater inter-subject variance and the lower effect sizes in P1 latency found in SCA-HSAC results could be explained by the removal of such negativity.

Regarding N2 latency, contrasting findings were provided by different approaches. In the original waveform, the AEP latency to piano sounds was significantly shorter than that to violin, whereas they were significantly shorter only to flute sounds in both SCA waveforms. The differences in latency between responses to piano and flute sounds after SCA might reflect the removal of interfering components that were masking N2 in piano responses, as flute latency value was not affected by SCA decomposition. Similarly, the difference between piano and violin found in the original waveform might have been caused by the overlapping of an earlier component that affected piano latency and that was removed after SCA. In addition, flute latency was shorter than violin latency in the SCA-TM waveform. However, as the violin condition provided the least successful results in SCA-TM (21/33 matches), this effect might be solely due to the lower number of subjects considered.

The most prominent difference in respect to the spectrotemporal properties of the stimuli was represented by the responses to the piano tones, which were consistent across all three methods. Compared to the other conditions, P1 component for the piano sounds had larger amplitude and shorter latency, whereas N2 amplitude was smaller. Furthermore, the N1-like component had its greatest amplitude for piano tones, which highlighted the presence of N1 already in the original waveform.

P1 and N2 amplitudes and latencies are known to decline with age during development, whereas N1 and P2 amplitudes gradually increase, to eventually become the dominant AEP components in adulthood (Ponton et al., 2000; Shahin et al., 2003; Tremblay et al., 2014). Changes in AEP morphology are thought to reflect cortical maturation processes such as changes in myelination (Albrecht et al., 2000; Caviness et al., 1996; Snook et al., 2005; Tonnquist-Uhlén et al., 1995) and cortical folding (Moore & Guan, 2001), favoring the selection of the most efficient networks for processing the information (Čeponiene et al., 1998).

In this regard, it has been suggested that changes in P1 and N2 features in early life might reflect differences in higher-level cognitive skills, playing a role comparable to that of P2 and N1 in adults (Čeponiene et al., 2002; Johnstone et al., 1996). For instance, enhanced P1 amplitude was found in children that received musical training as an index of experience-dependent plasticity (Habibi et al., 2016; Shahin et al., 2004), whereas previous studies in adults have linked enhanced N1/P2 to timbre discrimination (Jones et al., 1998; Meyer et al., 2006) and experience-dependent plasticity (Shahin et al., 2003; Tremblay et al., 2014) in adults. Our findings indicated the most prominent response to piano sounds, compared to the other two conditions. The auditory stimulation studied here was presented after another paradigm (musical multifeature or MuMufe), aiming to evoke mismatch negativity (MMN). The MuMufe involved the presentation of standard stimuli played on the piano, interleaved by deviant tones differing by six spectral properties (flute, violin, mistuning, omission, slide, and intensity). Notwithstanding, all three timbres – piano, flute, and violin – were present in this paradigm. However, piano sounds were presented 21 times more frequently than either violin or flute sounds. We, therefore, hypothesized that the reduced P1 and enhanced N2 amplitude for the piano condition that we observed in our results is a short-term plasticity effect, following the repeated presentation of piano sounds. We cannot rule out, however, that this observation could be an effect of an overall familiarity of our subjects with the sound of a piano. Habibi et al. (2016) described more prominent group differences between musically trained and non-trained children in their ERP responses to piano tones compared to those to violin and pure tones, despite the violin was the instrument the children played. Habibi and colleagues hypothesized that this could be due to the children having greater exposure to the sound of the piano since in their training program it was used in various music activities, e.g., for teaching music theory, altogether taking place more often than instrumental training. At the time of recording, subjects in our study did not follow any special musical training, though, they may have been exposed to the piano during regular musical activities in the school or kindergarten.

According to our initial hypotheses, SCA-HSAC demonstrated to provide reliable results at the individual level, compared to the conventional analysis approaches and the SCA-TM procedure. Moreover, differences in latencies following SCA reflected the separation of overlapping neural signals, providing a more reliable estimate of the true peaking latency of single responses. This allowed highlighting differences in responses to distinct sound features that were not visible in the waveforms analyzed with standard methods and eliminate spurious significances. We, therefore, propose that the ability to model neural signals at the individual level, together with the property of extracting the most consistent components, make of SCA-HSAC a promising tool for the use of ERPs in clinical settings.

## Acknowledgments

We wish to thank OrkesterMester program and its participants for collaboration on this project. We would like to thank Pætur Zachariasson and Sarah Foss for their assistance during the data collection. We thank Bjørn Petersen for his help with the sound stimuli. Center for Music in the Brain is funded by the Danish National Research Foundation (DNRF117).

